# Exploring new Bacteroidota strains: Functional Diversity and Probiotic Characteristics

**DOI:** 10.1101/2024.12.18.628894

**Authors:** Lisa Ladewig, Muhammad Aammar Tufail, Birhanu M. Kinfu, Hanna Fokt, John F. Baines, Ruth A. Schmitz

## Abstract

Bacteroidota, a diverse phylum of bacteria, are increasingly recognized for their significant contributions to host health, particularly through their antimicrobial and probiotic properties. This study investigates the functional diversity and probiotic potential of 42 new Bacteroidota strains enriched and identified from diverse hosts, including mouse ceca and human stool samples. Using 16S rRNA gene sequencing, we phylogenetically characterized the strains of the genera Bacteroides, Phocaeicola and Sphingobacterium and assessed their functional properties related to probiotic potential. The strains were evaluated concerning their ability to inhibit biofilm formation of WHO declared clinically significant pathogens, including gram-positive *Staphylococcus aureus* and *Staphylococcus epidermidis*, gram-negative *Klebsiella oxytoca* and *Pseudomonas aeruginosa*, and the eukaryotic fungus *Candida albicans.* Additionally, we investigated bile salt hydrolase and quorum quenching activities of the strains, key traits associated with probiotic efficacy. Our findings demonstrate that all examined Bacteroidota strains consistently exhibit a capacity to inhibit biofilm formation but to different extent. Furthermore, 14 strains showed quorum quenching activity, and 39 bile salt hydrolase activity, highlighting their probiotic potential. High biofilm inhibition as well as quorum quenching activity against both autoinducers, AHL and AI-2, were predominantly observed in *Bacteroides caecimuris* and *Bacteroides muris*, making them attractive candidates for next-generation probiotics. Overall, this study advances the field of next-generation probiotics by identifying promising candidates for therapeutic applications potentially revolutionizing approaches to microbiome-based interventions and pathogen control in clinical settings.

## 1. Introduction

In recent years, probiotic effects of several microorganisms have been discovered. Probiotics are live microorganisms, typically bacteria or yeasts, that are defined by the World Health Organization and the Food and Agriculture Organization as “live microorganisms that, when administered in adequate amounts, confer a health benefit on the host” (Latif et al., 2023). Traditional probiotics include bacterial species that belong to the genera *Lactobacillus* and *Bifidobacterium* (Latif et al., 2023). However, *Escherichia coli* Nissle 1917 (EcN), a Gram-negative Bacillus, has also emerged as a highly effective probiotic due to its colonization capabilities in the intestine and its ability to exhibit antagonistic effects against various pathogens. Additionally, EcN enhances the host’s immune function, making it one of the most widely used probiotics for the prevention and treatment of inflammatory bowel disease (IBD) (Zhao et al., 2022). These microorganisms function by competing with pathogens, maintaining an acidic pH, or producing molecules that support gut and immune functions. They also strengthen the gut barrier and have been linked to benefits such as reducing cholesterol levels, anti-carcinogenic activities, and lowering inflammation (recently reviewed in (Stavropoulou & Bezirtzoglou, 2020; Tufail & Schmitz, 2024). By reducing infections caused by pathogenic microorganisms, probiotics can potentially decrease the need for antibiotics, helping combat antibiotic resistance. Thus, probiotics are essential for maintaining a balanced gut microbiota, which is critical in preventing diseases.

In response to the need for more targeted and effective treatments, research has increasingly focused on Next-Generation Probiotics (NGPs), a new category of probiotic microorganisms. NGPs differ from traditional probiotics in that they are specialized to target particular diseases and are designed to act on precise biological mechanisms (Chang et al., 2019). These microorganisms are integral to the gut microbiota and have the potential to enhance the immune system, strengthen the gut barrier, and positively influence metabolism (Hasnain et al., 2024).

Most probiotics are isolated from food sources or the human gut microbiota itself (Al-Fakhrany & Elekhnawy, 2024). The gut microbiota of other hosts also offers a suitable source for probiotics. The hunt for probiotics from these hosts holds promise for discovering new NGP strains with unique health benefits (Fontana et al., 2013; Venkatesh et al., 2024). However, this field remains underdeveloped particularly due to safety concerns, regulatory barriers, and a focus on established sources like human gut microbiota and dairy products (Venkatesh et al., 2024).

Despite these obstacles, research on animal-derived NGPs is crucial, as it might reveal bacterial strains with better resistance against pathogens and improved microbiome diversity, enhancing therapeutic options for gut health. There are two main strategies for developing NGPs: either linking a specific strain to a health benefit or using a well-characterized probiotic strain to deliver beneficial molecules that address disease and promote health (O’Toole et al., 2017). Isolates from the phylum Bacteroidota, previously known as Bacteroidetes, are among the promising candidates for next-generation probiotics (Tan et al., 2019). This phylum encompasses a diverse group of Gram-negative, obligately anaerobic bacteria that are ubiquitous in the human gut microbiota (Karlsson et al., 2011).

Recent studies have highlighted the potential of Bacteroidota strains, such as *Bacteroides fragilis* and *B. thetaiotaomicron*, in modulating host immune responses and ameliorating various diseases (He et al., 2023; Lalowski & Zielińska, 2024; Cang et al., 2024; Qu et al., 2022). In our recent review, we summarized the studies highlighting the potential of *Bacteroides* spp. as next-generation probiotics, focusing on their ability to maintain gut health and modulate immune responses and highlighted the promising therapeutic applications of *Bacteroides* spp., while also acknowledging the necessity for safety assessments due to their opportunistic pathogenic nature (Tufail & Schmitz, 2024). In addition to the probiotic properties, Bacteroidota has the ability to adapt to the gut environment and interact with host tissues makes Bacteroidota an ideal candidate for next-generation probiotics (Tan et al., 2019). Building upon these promising attributes of Bacteroidota strains, a crucial aspect in evaluating their potential as NGPs is their antimicrobial activity profile. While the above-mentioned studies demonstrate their immunomodulatory and metabolic benefits, understanding their direct antagonistic effects against pathogens is equally important. This includes assessing not only their ability to inhibit pathogen growth, but also their capacity to disrupt biofilm formation - a key virulence mechanism that shields pathogenic microorganisms from both antimicrobial agents and host immune responses. The ability to prevent or disrupt biofilms could complement the already established beneficial properties of Bacteroidota strains, potentially enhancing their therapeutic efficacy as NGPs.

Probiotic effects are achieved by competing with pathogens for nutrients and attachment sites, producing antimicrobial substances, and altering environmental conditions, such as lowering pH levels (Vuotto et al., 2014). Another important mechanism is quorum quenching, where probiotics disrupt bacterial communication processes that regulate biofilm formation and virulence (Ghanei-Motlagh et al., 2020). Additionally, bile salt hydrolysis is important for probiotic bacteria since it helps them to survive in the gut, regulate the microbiome and reduce the toxic effects of bile salts, thus enabling them to establish themselves and exert their positive effects (Mohanty et al., 2024). This makes probiotics valuable in combating infections, especially those caused by antibiotic-resistant bacteria (Ghanei-Motlagh et al., 2020).

The aim of this study was to increase the diversity of *Bacteroides* isolates by obtaining new strains from diverse host samples and conduct a comprehensive assessment of their potential as NGPs. Initial 16S rRNA gene sequencing was used for taxonomical identification and classification. This genetic characterization provided a solid foundation for understanding the diversity and phylogenetic relationships of the new isolated strains. Following we performed a series of tests to assess the strains’ functional properties. These included high-throughput biofilm inhibition screening, microfluidic flow cell visualization of biofilm inhibition, quorum quenching assessment and bile salt hydrolase activity analysis. In particular, there is a gap in our current knowledge regarding the QQ activities of Bacteroidota, which emphasize the significance of our discovery of QQ activity of 14 isolates. Overall, the significance of this multifaceted approach lies in its ability to provide a holistic understanding of the strains’ capabilities and their potential applications in promoting gut health and combating pathogenic microorganisms.

## 2. Materials and Methods

### 2.1. Enrichment, Isolation and characterization of Bacteroidota strains

Samples were collected from mouse cecal contents (*Mus Musculus*) (Max Planck Institute for Evolutionary Biology, Plön, Germany), feces from tortoise, horse, cow, guinea pig, rabbit, muoflon, hedgehog, galloway, cat, goose (Kiel, Germany) and human stool from a healthy donor (Kiel, Germany) as shown in Table 1. Mice were maintained and handled according to FELASA guidelines and German animal welfare law (Tierschutzgesetz § 11, permit from Veterinäramt Kreis Plön: 1401–144/PLÖ–004697). Sampling of human stool was conducted by Dr. Corinna Bang in accordance with the Declaration of Helsinki. Ethical approval was granted by the ethics committee at Kiel University (D590/22). All volunteers provided written informed consent.

**Table 1:** Taxonomic identification of isolated Bacteroidota strains.

| Identification<br>by 16S rRNA<br>gene analysis | Isolate | Source<br>sample | Best match | Scientific<br>name | Accession<br>No. | Query<br>Coverage<br>(%) | Identity<br>(%) |
| --- | --- | --- | --- | --- | --- | --- | --- |
| *Bacteroides*<br>*caecimuris* | 241174E | Mouse<br>cecum | Bacteroides<br>caecimuris strain<br>I48 chromosome | *Bacteroides*<br>*caecimuris* | CP015401.2 | 99 | 99.64 |
|  | 241175E | Mouse<br>cecum | Bacteroides<br>caecimuris strain<br>NM63_1-25 16S<br>ribosomal RNA<br>gene | *Bacteroides*<br>*caecimuris* | MK929076.<br>1 | 99 | 99.58 |
|  | 241176E | Mouse<br>cecum | Bacteroides<br>caecimuris strain<br>NM63_1-25 16S<br>ribosomal RNA<br>gene | *Bacteroides*<br>*caecimuris* | MK929076.<br>1 | 100 | 99.57 |
|  | 241160E | Mouse<br>cecum | Bacteroides<br>caecimuris strain<br>0.1XD8-16 16S<br>ribosomal RNA<br>gene | *Bacteroides*<br>*caecimuris* | MK287726.<br>1 | 99 | 99.49 |
|  | 241177E | Mouse<br>cecum | Bacteroides<br>caecimuris strain<br>NM63_1-25 16S<br>ribosomal RNA<br>gene | *Bacteroides*<br>*caecimuris* | MK929076.<br>1 | 100 | 99.65 |
|  | 241178E | Mouse<br>cecum | Bacteroides<br>caecimuris strain<br>I48 chromosome | *Bacteroides*<br>*caecimuris* | CP015401.2 | 98 | 99.57 |
|  | 241159E | Mouse<br>cecum | Bacteroides<br>caecimuris strain<br>0.1XD8-16 16S<br>ribosomal RNA<br>gene | *Bacteroides*<br>*caecimuris* | MK287726.<br>1 | 100 | 99.28 |
| ***Bacteroides***<br>***fragilis*** | 241179E | Human<br>stool | Bacteroides<br>fragilis strain<br>GUT04<br>chromosome | *Bacteroides*<br>*fragilis* | CP043610.1 | 99 | 98.21 |
|  | 241171E | Human<br>stool | Bacteroides<br>fragilis strain<br>BFG-7<br>chromosome | *Bacteroides*<br>*fragilis* | CP103244.1 | 99 | 99.72 |
|  | 241180E | Human<br>stool | Bacteroides<br>fragilis strain<br>GUT04<br>chromosome | *Bacteroides*<br>*fragilis* | CP043610.1 | 99 | 99.75 |
|  | 241172E | Human<br>stool | Bacteroides<br>fragilis strain<br>GUT04<br>chromosome | *Bacteroides*<br>*fragilis* | CP043610.1 | 100 | 99.67 |
|  | 241181E | Human<br>stool | Bacteroides<br>fragilis strain<br>GUT04<br>chromosome | *Bacteroides*<br>*fragilis* | CP043610.1 | 100 | 99.72 |
|  | 241182E | Human<br>stool | Bacteroides<br>fragilis strain P207<br>chromosome | *Bacteroides*<br>*fragilis* | CP114371.1 | 94 | 90.66 |
|  | 241170E | Human<br>stool | Bacteroides<br>fragilis strain<br>GUT04<br>chromosome | *Bacteroides*<br>*fragilis* | CP043610.1 | 100 | 99.83 |
|  | 241169E | Human stool | Bacteroides fragilis strain GUT04 chromosome | *Bacteroides fragilis* | CP043610.1 | 99 | 100 |
|  | 241183E | Horse feces | Bacteroides fragilis strain GUT04 chromosome | *Bacteroides fragilis* | CP043610.1 | 100 | 98.21 |
|  | 241173E | Tortise feces | Bacteroides fragilis strain GUT04 chromosome | *Bacteroides fragilis* | CP043610.1 | 99 | 99.91 |
|  | 241188E | Human stool | Bacteroides fragilis isolate 12-13--1-44 16S ribosomal RNA gene, partial sequence | *Bacteroides fragilis* | PP465593. | 100 | 100 |
| ***Bacteroides muris*** | 241155E | Mouse cecum | Bacteroides muris KH365_2 16S ribosomal RNA gene | *Bacteroides muris* | ON325392.1 | 100 | 97.91 |
|  | 241156E | Mouse cecum | Bacteroides muris KH365_2 16S ribosomal RNA gene | *Bacteroides muris* | ON325392.1 | 100 | 97.91 |
| ***Bacteroides ovatus*** | 241158E | Human stool | Bacteroides ovatus partial 16S rRNA gene, strain 23736 | *Bacteroides ovatus* | LR999666.1 | 100 | 100 |
|  | 241184E | Human stool | Bacteroides ovatus strain CP926 chromosome | *Bacteroides ovatus* | CP113514.1 | 99 | 99.37 |
| *Bacteroides xylanisolvens* | 241185E | Human stool | Bacteroides xylanisolvens strain H207 chromosome | *Bacteroides xylanisolve*<br>*ns* | CP041230.1 | 99 | 95.95 |
|  | 241167E | Human stool | Bacteroides xylanisolvens strain H207 chromosome | *Bacteroides xylanisolve*<br>*ns* | CP041230.1 | 98 | 98.02 |
|  | 241165E | Human stool | Bacteroides xylanisolvens strain APCS1/XY chromosome | *Bacteroides xylanisolve*<br>*ns* | CP042282.1 | 99 | 99.74 |
|  | 241186E | Human stool | MAG: Bacteroides xylanisolvens isolate KR001_HAM_00 12 chromosome | *Bacteroides xylanisolve*<br>*ns* | CP107195.1 | 99 | 99.91 |
|  | 241168E | Human stool | Bacteroides xylanisolvens strain AY11-1 isolate human Intestinal chromosome | *Bacteroides xylanisolve*<br>*ns* | CP120351.1 | 99 | 99.79 |
|  | 241187E | Human stool | Bacteroides xylanisolvens strain AY11-1 isolate human Intestinal chromosome | *Bacteroides xylanisolve*<br>*ns* | CP120351.1 | 99 | 99.51 |
|  | 241189E | Human stool | Bacteroides xylanisolvens strain H207 chromosome | *Bacteroides xylanisolve*<br>*ns* | CP041230.1 | 100 | 98.73 |
|  | 241166E | Human stool | Bacteroides xylanisolvens strain AY11-1 isolate human Intestinal chromosome | *Bacteroides xylanisolve ns* | CP120351.1 | 100 | 99.50 |
|  | 241190E | Human stool | Bacteroides xylanisolvens strain APCS1/XY chromosome | *Bacteroides xylanisolve ns* | CP042282.1 | 99 | 97.98 |
| ***Phocaeicola sartorii*** | 241161E | Mouse cecum | Phocaeicola sartorii strain DSM 110081 16S ribosomal RNA gene | *Phocaeicol a sartorii* | OM658617.1 | 100 | 98.95 |
|  | 241162E | Mouse cecum | Phocaeicola sartorii strain DSM 110081 16S ribosomal RNA gene | *Phocaeicol a sartorii* | OM658617.1 | 99 | 99.4 |
|  | 241191E | Mouse cecum | Phocaeicola sartorii strain DSM 110081 16S ribosomal RNA gene | *Phocaeicol a sartorii* | OM658617.1 | 100 | 99.51 |
|  | 241192E | Mouse cecum | Phocaeicola sartorii strain DSM 110081 16S ribosomal RNA gene | *Phocaeicol a sartorii* | OM658617.1 | 99 | 99.51 |
|  | 241163E | Mouse<br>cecum | Phocaeicola<br>sartorii strain DSM<br>110081 16S<br>ribosomal RNA<br>gene | *Phocaeicol*<br>*a sartorii* | OM658617.<br>1 | 100 | 99.44 |
|  | 241193E | Mouse<br>cecum | Phocaeicola<br>sartorii strain DSM<br>110081 16S<br>ribosomal RNA<br>gene | *Phocaeicol*<br>*a sartorii* | OM658617.<br>1 | 100 | 99.51 |
|  | 241194E | Mouse<br>cecum | Phocaeicola<br>sartorii strain DSM<br>110081 16S<br>ribosomal RNA<br>gene | *Phocaeicol*<br>*a sartorii* | OM658617.<br>1 | 99 | 99.02 |
|  | 241164E | Mouse<br>cecum | Phocaeicola<br>sartorii strain DSM<br>110081 16S<br>ribosomal RNA<br>gene | *Phocaeicol*<br>*a sartorii* | OM658617.<br>1 | 99 | 99.02 |
|  | 241195E | Mouse<br>cecum | Phocaeicola<br>sartorii strain DSM<br>110081 16S<br>ribosomal RNA<br>gene | *Phocaeicol*<br>*a sartorii* | OM658617.<br>1 | 99 | 99.23 |
| ***Sphingobacter***<br>***ium***<br>***multivorum*** | 241196E | Tortise<br>feces | Sphingobacterium<br>multivorum strain<br>CSpm_LPR10+<br>16S ribosomal<br>RNA gene, partial<br>sequence | *Sphingobac*<br>*terium*<br>*multivorum* | MH788992.<br>1 | 99 | 99.51 |
|  | 241197E | Tortise<br>feces | Sphingobacterium<br>multivorum strain<br>CSpm_LPR10+<br>16S ribosomal<br>RNA gene, partial<br>sequence | *Sphingobac*<br>*terium*<br>*multivorum* | MH788992.<br>1 | 99 | 99.65 |

The samples were serially diluted in 1 x PBS buffer (137 mM NaCl, 2.7 mM KCI, 10 mM Na2HPO4, 1.8 mM KH2PO4, pH 7.4) in an anaerobic chamber. The strains were anaerobically isolated by plating the diluted samples on Schaedler anaerobe KV selective agar with lysed horse blood (Thermo Fisher Scientific, Darmstadt, Germany) as well as on a non-selective Schaedler anaerobe agar with sheep blood (Thermo Fisher Scientific, Darmstadt, Germany), Columbia agar (Carl Roth, Karlsruhe, Germany), and anaerobe Basal agar (Thermo Fisher Scientific, Darmstadt, Germany). Incubation was done at 37 °C under anaerobic conditions.

Colonies were re-streaked for purification onto the respective agar plates for three times and the single colonies were inoculated in 5 mL pre-reduced modified chopped meat broth (20 g/L meet extract, 30 g/L casein peptone, 5 g/L yeast extract, 5 g/L K2HPO4, 1 g/L glucose, 5 mg/L, 0.1 mg/L, L-cysteine 0.5 g/L, Haemin 10 mL/L Vitamin K1 10 mL/L) and grown anaerobically at 37 °C in closed Hungate tubes. Cells were cryopreserved with ROTI®Store cryo beads (Carl Roth, Karlsruhe, Germany) and stored at −80 °C.

Initial study of Bacteroidota strains was done with colony check PCR using the genus-specific primer set Primer_Bac_Fw (5′-GGCGCACGGGTGAGTAAC-3′) (Gómez-Doñate et al., 2016) and AllBAc412r (5′-CGCTACTTGGCTGGTTCAG-3′) (Layton et al., 2006) as well as phylum-specific primer pairs Bac1100R (5′-TTAASCCGACACCTCACGG-3′) (Yang et al., 2015) and S-P-Bdet-0107-a-S-21(5′-GCACGGGTGAGTAACACGTA-3′) (Pfeiffer et al., 2014). PCR amplification was performed using purified DNA as template and GoTaq Polymerase (Promega, Walldorf, Germany) with the following protocol.95 °C, 2 min followed by 35 cycles (95°C, 30 sec; 55 °C, 30 s; 72 °C, 90 s), 5 min extension at 72 °C. To generate full-length 16SrRNA genes universal 16SrRNA primer 16S_27F (5′-AGAGTTTGATCCTGGCTCAG-3′) and 16S_1492R (5′-CGGTTACCTTGTTACGACTT-3′) (Petruska et al., 1988) were used. In addition, an amplified ribosomal DNA restriction analysis (ARDRA) was performedto distinguish between different strains based on the restriction patterns. Therefore, amplified 16S rRNA genes were digested with *BamH*I*, Nhe*I*, Nae*I*, Pst*I*, EcoR*I and *Dra*I in CutSmart Buffer (NEB, Frankfurt am Main, Germany) to completeness digestion and analyzed on agarose gel. For further taxonomic identification, patterns were grouped, and representatives were selected for full 16S rRNA sequencing (Institute of Clinical Molecular Biology, Kiel, Germany). The 16S rRNA sequence data were submitted to GenBank under accession numbers PQ775146-PQ775187. The variability between Bacteroidota strains that were very similar in their 16S rRNA sequences was further checked using Random Amplified Polymorphic DNA (RAPD)-PCR to select representative strains. Therefore, the primer set P1 (5′-CCGCAGCCAA-3′) and P2 (5′-AACGGGCAG-3′) (Gutiérrez et al., 2011) was used. Amplification was performed using GoTaq Polymerase with an initial denaturation at 94 °C for 3 minutes, followed by 40 cycles of denaturation at 94 °C for 45 seconds, annealing at 35 °C for 30 seconds, and extension at 72 °C for 1 minute, with a final extension step at 72 °C for 10 minutes. Sequences were edited and analyzed using Geneious Prime 2022.2. 2 (Dotmatics, Bishop’s Stortford, UK).

For phylogenetic tree construction, the evolutionary history was inferred using the Neighbor-Joining method (Saitou & Nei, 1987). The bootstrap consensus tree inferred from 1000 replicates (Felsenstein, 1985) is taken to represent the evolutionary history of the taxa analyzed (Saitou & Nei, 1987). Branches corresponding to partitions reproduced in less than 50% bootstrap replicates are collapsed. The percentage of replicate trees in which the associated taxa clustered together in the bootstrap test (1000 replicates) are shown below the branches (Saitou & Nei, 1987). The evolutionary distances were computed using the Maximum Composite Likelihood method (Tamura et al., 2004) and are in the units of the number of base substitutions per site. This analysis involved 42 nucleotide sequences. All ambiguous positions were removed for each sequence pair (pairwise deletion option). There were a total of 3440 positions in the final dataset. Phylogenetic tree of evolutionary relationship of taxa among our strains and selected type strains was generated using MEGA11 (Stecher et al., 2020; Tamura et al., 2021).

### 2.2. Preparation of cell-free supernatant, cell extract, and subcellular fractions from Bacteroidota strains for biofilm analysis

The Bacteroidota strains were cultured in modified chopped meat broth at 37 °C overnight. A total culture volume of 50 mL in closed serum bottles was utilized for both the static and dynamic biofilm inhibition assays. Exponential growing cultures were centrifuged at 4.000 x g and 4 °C for 1 h. After centrifugation, cell-free culture supernatant was additional filtered through 0.2 µm filters (Sarstedt, Nümbrecht, Germany). Cell pellet was re-suspended in 4 mL of 50 mM Tris/HCL buffer (pH 7.9) containing 50 mM NaCl, and transferred into screwcap vial (1 ml? Sarstedt, Nürnberg, Germany) containing 0.1 mm and 2.5 mm glass beads (Carl Roth, Karlsruhe, Germany). Cells were mechanically disrupted using Precellys 24 homogenizer (Bertin Technologies, Montigny-le-Bretonneux, France) at 5000 x g for 3 × 30 s. Samples were centrifuged at 15.000 x g and 4 °C for 30 min. The clear supernatant was filtered through 0.2 µm spin filter units (Amchro, Hattersheim, Germany) generating the soluble cell extract. The obtained cell extract was subsequently filtered through 10 kDa followed by a 3 kDacentrifugal filter (Merck KGaA, Darmstadt, Germany) to generate three sub-cellular fractions based on molecular size (> 10 kDa, 3 to 10 kDa, and < 3 kDa fractions).

### 2.3. Evaluating quorum quenching activities in Bacteriodota strains

The cell extract and culture supernatant of Bacteriodota strains grown under anaerobic conditions in 5 mL cultures in closed Hungate tubes were generated (see above) and screened for interference with acylated homoserine lactone (AHL) and autoinducer 2 (AI2) so called quorum quenching activities - using two reporter strains, AI1-QQ.1 and AI2-QQ.1 (Weiland-Bräuer et al., 2015). QQ activity was determined using the methodology reported in detail by Weiland-Bräuer et al., (2015). The Assay was performed with three independent biological replicates with each four technical replicates

### 2.4. Inhibition of pathogenic biofilm formation using static biofilm assay

The pathogenic biofilm forming strains *Klebsiella oxytoca* M5aI (DSM 7342), *Candida albicans* (DSM 11225), *Staphylococcus aureus* (DSM 11323), *Staphylococcus epidermidis* RP62A (DSM 28319) and *Pseudomonas aeruginosa* PAO1 (DSM 1707) were used to screen for inhibition activities of Bacteroidota straines*C. albicans* was grown overnight in 5 mL YPD (20 g/L glucose, 20 g/L peptone, 10 g/L yeast extract) broth while all others were grown in Luria Bertani broth (LB, Carl Roth, Karlsruhe, Germany) shaking at 120 rpm. *C. albicans* and *K. oxytoca* M5al cultures were incubated at 30 °C and the remaining at 37 °C. Upon completion of overnight cultivation, Neubauer cell counting was employed to analyze the cell concentrations, which were subsequently adjusted to 1 × 10^6^ cells/mL using GC minimal broth (with 1% (v/v) glycerol and 0,3% (w/v) casmino acids) for *K. oxytoca* M5aI, YPD broth for *C. albicans* and Caso broth (17 g/L casein peptone, 3 g/L soybean peptone, 5 g/L NaCl, 2.5 g/L K2HPO4, and 2.5 g/L glucose) for the remaining strains. Following, the cultures were dispensed into a 96-well round (U)-bottom microtiter plate (Greiner Bio-One, Kremsmünster, Austria), with each well containing 190 µL. To each well, 10 μL of culture filtered supernatant, cell-free supernatant or subcellular fractions derived from Bacteroidota strains was supplemented. This was performed using three biological replicates with each eight technical replicates. Tris-HCl buffer (50 mM, pH 7.9) was used as a control. The microtiter plates were sealed with adhesive gas-permeable sterile membrane (Breathe Easy, Carl Roth, Karlsruhe, Germany) and incubated at 18 h at the appropriate temperatures.

The biofilm assay was conducted utilizing the Crystal Violet assay (Haney et al., 2021), employing a newly devised 3D-printed wash assist device (Fig. S1). The design was crafted using Autodesk Inventor software (Munich, Germany) and Prusa Slicer (Prague, Czech Republic). Specifically engineered pins were incorporated to facilitate the unimpeded flow of liquid from wells, minimizing contact with the walls and thus reducing disturbance to the biofilm. Pin apertures aid in air circulation and facilitate the rupture of liquid surface tension. The final design was 3D-printed using polyethylene terephthalate glycol on a Prusa MK3S printer (Technical facility, Christian-Albrechts-Universität zu Kiel, Kiel, Germany).

The microtiter plate was fitted upside-down to the wash assist device allowing gentle flow of culture medium containing planktonic cells from the wells. The biofilm was washed two times with 200 μL/well 1 x PBS buffer (137 mM NaCl, 2.7 mM KCI, 10 mM Na2HPO4, 1.8 mM KH2PO4, pH 7.4) using the wash assist device, described above. Intact biofilm attached to the wells was stained by adding 200 μL/well of 0.1 % (w/v) crystal violet solution (Carl Roth, Karlsruhe, Germany) and incubated at room temperature for 30 min. The crystal violet solution was removed and the wells subsequently washed two times with 1 x PBS buffer as described above. After drying, the stained biofilm was extracted by adding 200 μL/well of 80% ethanol and shaking at 80 rpm for 30 min at room temperature to extract the crystal violet. The absorbance of the resolved crystal violet, a quantitative indicator of biofilm formation, was monitored at 590 nm using a plate spectrophotometer (Spectra max Plus 384, Molecular Devices, Ismaning, Germany), the inhibitory activity was calculated relative to the respective control.

### 2.5. Biofilm inhibition in a microfluidic flow cell system using Bacteroidota strains

The biofilm formation was performed for both *K. oxytoca* M5al and *S. epidermidis* RP62A. The methodology of analyzing biofilm formation in a costume made microfluidic flow cell (channel volume of 0.8 µL µL) has been established and descripted in Ladewig et al. (2023) for *K. oxytoca* M5al. For *S. epidermidis* RP62A, the conditions had to be newly established. After optimization the biofilm formation was conducted in Caso broth at an incubation temperature of 37 °C for 20 h. Inoculation was performed using a cell suspension of *S. epidermidis* RP62A of 4 ×10^8^ cells/mL. For biofilm formation, initial cell adhesion was ensured by incubation for 1 h without any flow. After that, the channel was continuously rinsed through the first inlet at a defined flow rate. For *K. oxytoca* M5al, a flow rate of 15 µL/h, and for *S. epidermidis* RP62A, a flow rate of 18 µL/h was applied. To test inhibition effects of new isolates the respective medium was supplemented with cell free cell extract to a final concentration of 0.5%. The running time for *S. epidermidis* was 20 h and for *K. oxytoca* 24 h, with continuous flow. The cell extract-containing reservoir was renewed every 6 h during the running time. For *K. oxytoca* M5al the cell free cell extract of the strains 241155E and 241174E were used. For *S. epidermidis* RP62A the cell extract of the strains 241155E and 241170E were chosen. Biofilm formation was carried out at the corresponding incubation conditions and flow rate. As a control biofilm formation of *K. oxytoca* M5aI and *S. epidermidis* RP62A was studied without cell extract addition. Four biological replicates were conducted for each treatment.

The analysis of biofilm formation was conducted as described in Ladewig et al., (2023). Four each image stacks were captured with 0.9 µm z-steps, and biofilm parameters were calculated using “Zen Black” (version 14.0.22.201) (Zeiss) and “Imaris” (version 9.9.0) (Oxford Instruments) software.

### 2.6. Investigation of bile salt hydrolase activity of Bacteriodota strains

The BSH activity was analysed using the modified anaerobe basal broth (Thermo Fisher Scientific, Darmstadt, Germany). The anaerob basal broth was supplemented with 375 mg/L CaCl2 (Sigma-Aldrich Chemie GmbH, Taufkirchen, Germany) and 2.5 g/L porcine bile extract (Sigma-Aldrich Chemie GmbH, Taufkirchen, Germany) to modified anaerobic basal agar, containing 20 g/L agar. 20 mL of modified anaerobic basal agar was poured into round petri dishes and stored anaerobically in the dark without sunlight until needed. Agar plates were punched using the back of a sterile pipette tip to generate an 8 mm hole in the agar. Bacteroides preculture was growing in 300 μL AB broth and incubated for 72 hours at 37 °C. The main culture was grown in 300 μL AB broth for 24 hours at 37 °C. The OD600 was adjusted to 1.0 and 75 μL of the main culture was filled to the punched hole of AB agar and carefully incubated at 37 °C for 48 h under anaerobic conditions. This was done using three biological replicates. The diameter of precipitation zones around the punched holes was measured for activity comparison.

## 3. Results

### 3.1. Diversity and identification of Bacteroidota strains from animal and human samples

Initially, animal and human samples suspected to have a high abundance and diversity of Bacteroidota representatives were collected. These samples included animal fecal, mouse cecum, and human stool samples (Table 1). Through enrichment on anaerobic selective and non-selective media, a total of 690 morphologically distinct bacterial colonies were identified. The single colonies were further purified and screened via PCR using Bacteroidota-specific primers to exclude non-Bacteroidota species, followed by ADAR to eliminate redundant strains. Overall, 42 strains belonging to the Phylum Bacteroidota were identified (Table 1). Further sequence analysis of the full length 16S rRNA genes revealed a significant diversity within the 42 strains, identifying seven distinct species within this group including *Bacteroides caecimuris*, *Bacteroides fragilis*, *Bacteroides muris*, *Bacteroides ovatus*, *Bacteroides xylanisolvens*, *Phocaeicola sartorii*, and *Sphingobacterium multivorum* (Fig. 1). *B. caecimuris* and *P. sartorii* were the predominant species identified in mouse cecum samples, while *B. fragilis* and *B. xylanisolvens* were most frequently isolated from human samples.

**Figure 1:**
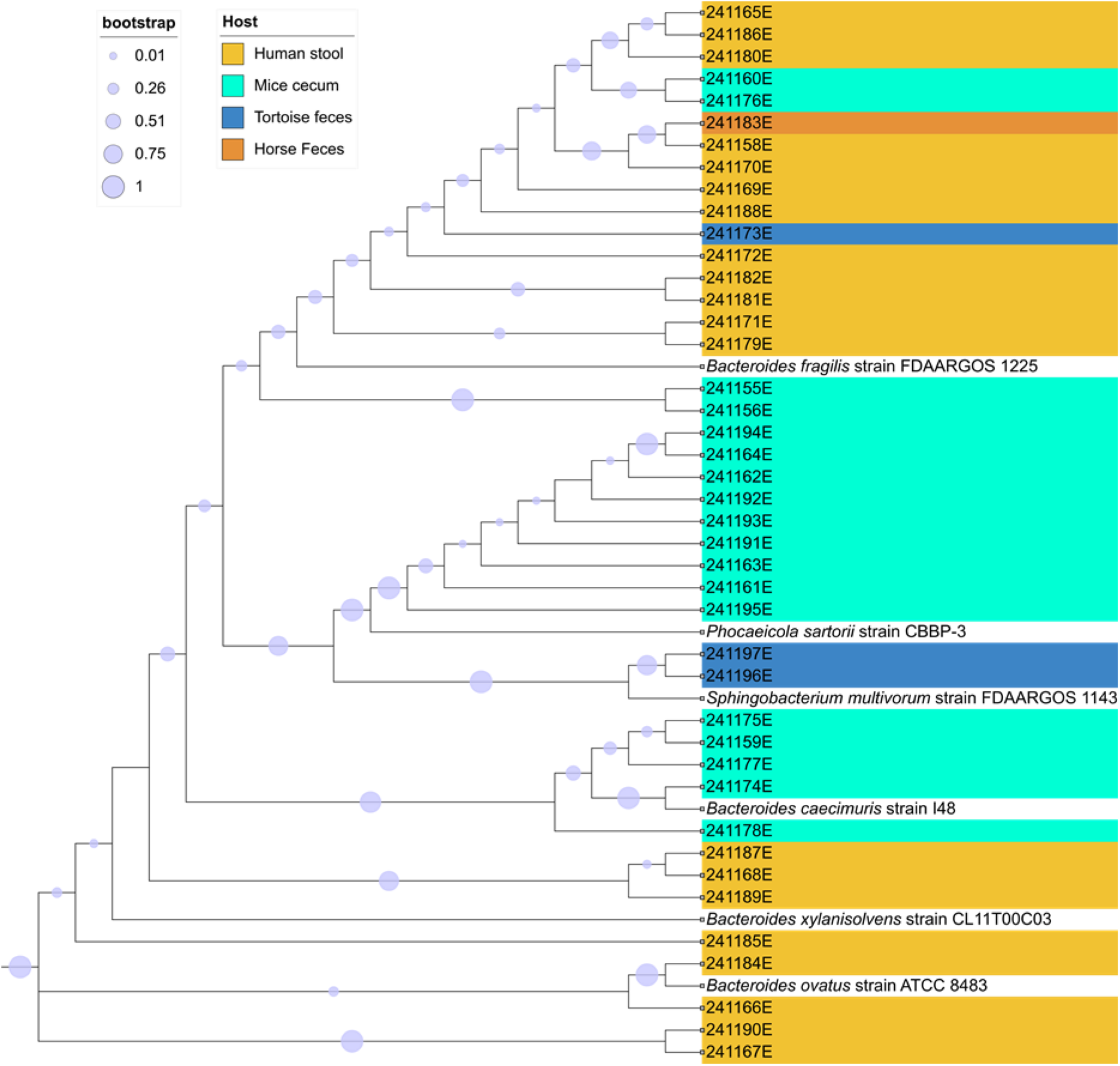
Phylogenetic tree illustrating the evolutionary relationships between the isolated strains (shaded in different colors based on the host name) and established type strains/reference genomes (unshaded) of Bacteroidota species. The tree depicts the genetic distances and relatedness among the taxa, highlighting the placement of our isolates within the broader phylogeny of Bacteroidota. Bootstrap values (where applicable) indicate the robustness of each clade, represented with size of circle, supporting the phylogenetic positioning of the isolates relative to known species (unshaded) of the Bacteroidota phylum.

### 3.2. The majority of the new bacteroidota strains show biofilm inhibiting activity

The 42 Bacteroidota strains were evaluated for their ability to inhibit biofilm formation of five opportunistic pathogens, including Gram-negative bacteria *K. oxytoca* M5aI and *P. aeruginosa* PA01, as well as Gram-positive bacteria *S. aureus* and *S. epidermidis*, and the yeast *C. albicans* known for causing fungal infections. Culture supernatants and three cellular fractions (> 10 kDa, 3-10 kDa, < 3 kDa), generated from exponentially growing Bacteroidota strains, were evaluated using a microtiter plate-based crystal violet assay as described in 2.4. In total, 168 samples were analyzed using three biological replicates each with eight technical replicates per sample. Reductions in biofilm formation were observed across various fractions for different pathogens as depicted in Figs. 2 and 3. 116 samples demonstrated a significant reduction in biofilm formation (≥ 20%) of at least one of the analyzed pathogens (Fig. 2). Notably, 34 culture supernatants and 36 fractions with a molecular size > 10 kDa were identified to be effective in inhibiting biofilm formation. Additionally, 23 samples of each smaller molecular size range (3-10 kDa and < 3 kDa) also showed an inhibition of biofilm formation (see Fig. 3). Whereas high numbers for strains inhibiting *S. epidermidis* and *S. aureus* as well as *K. oxytoca* were identified, significant less inhibiting fractions were identified for *P. aeroginosa* and *C. albicans*. In detail, *C. albicans* exhibited reductions up to 28% in six samples, while *K. oxytoca* showed reductions up to 50% across 33 samples, and *P. aeruginosa* demonstrated reductions up to 38% in seven samples. Furthermore, significant reductions up to 69 % were observed for *S. aureus* in 40 samples, and for *S. epidermidis*, with reductions up to 82 % in 82 samples as shown in Fig. 2. It is notably that most of the tested samples showed an inhibition against the Gram-positive pathogens *S. aureus* and *S. epidermidis*.

**Figure 2:**
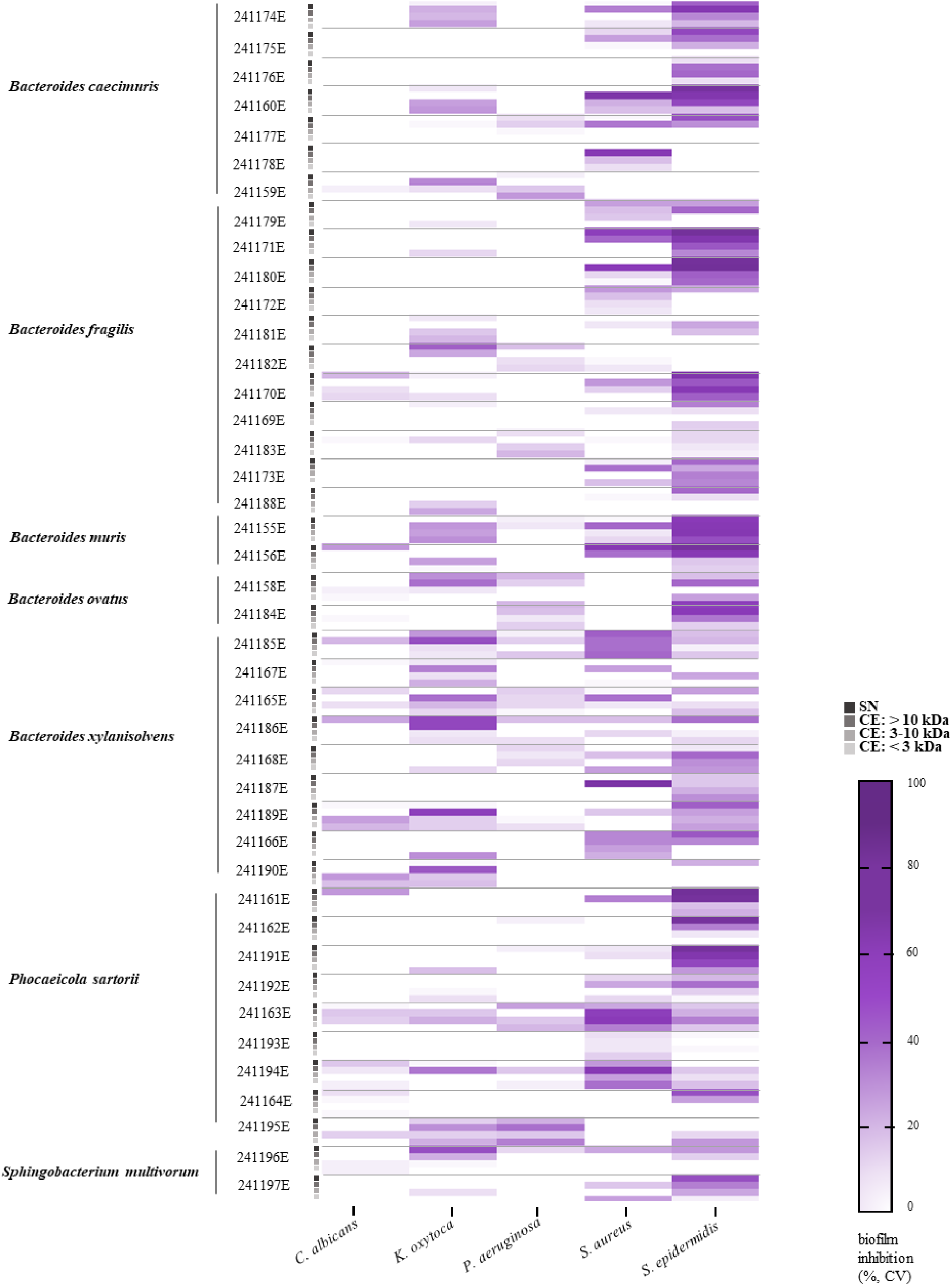
Inhibition of biofilm formation by Bacteroidota isolates. The heatmap illustrates the percentage of biofilm inhibition calculated relative to the corresponding biofilm control. Supernatant and three subcellular fractions (grey intensity) from the cell extract of Bacteroidota isolates were tested to prevent biofilm formation of *C. albicans, K. oxytoca*, *P. aeruginosa, S. aureus* and *S. epidermidis*. The evaluation was performed with three biological replicates, each consisting of eight technical replicates. The intensity of the purple tiles indicates the level of inhibition.

**Figure 3:**
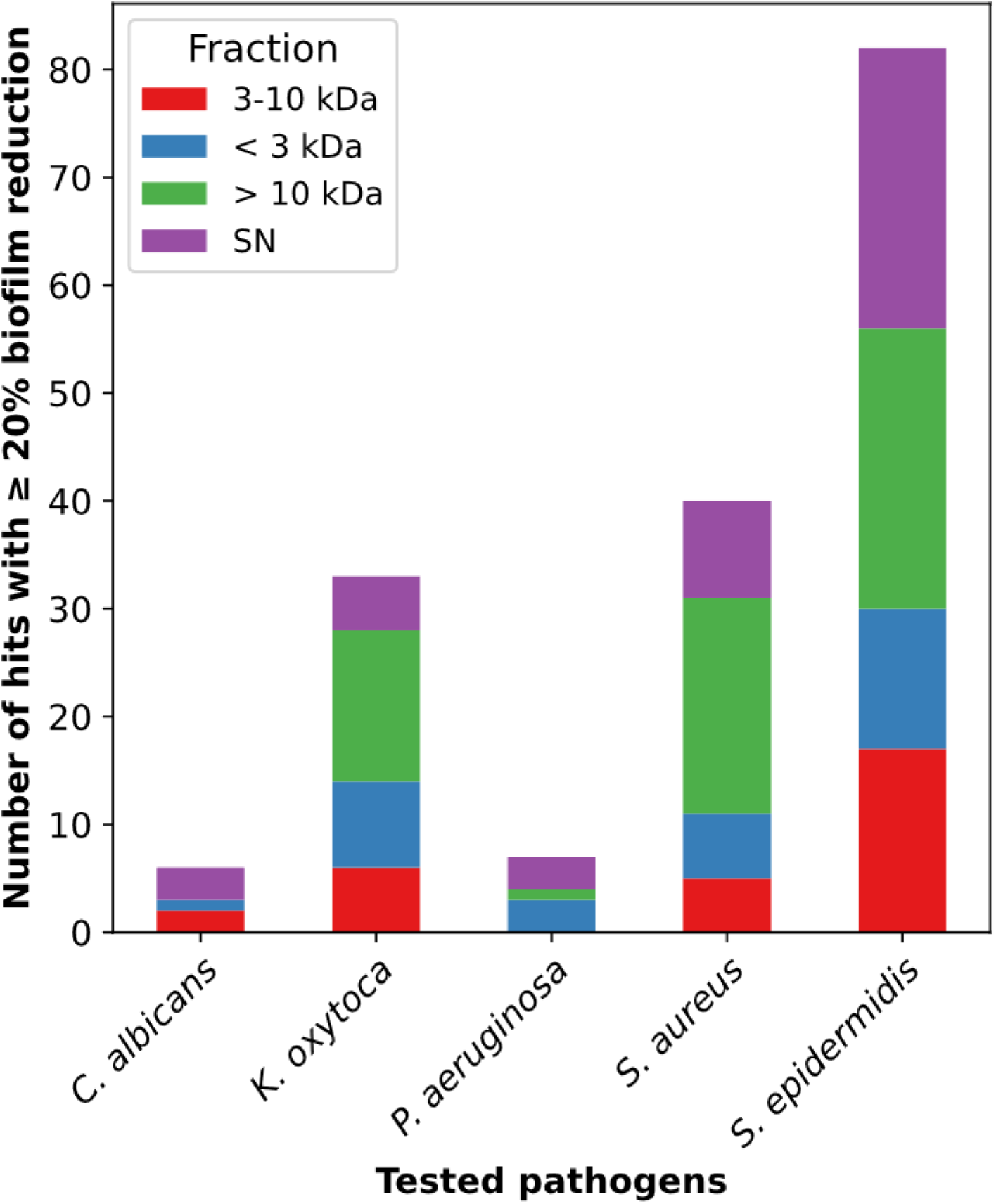
Distribution of biofilm reduction activity across tested pathogens. The graph shows the number of hits with ≥20% biofilm reduction for the pathogens *C. albicans, K. oxytoca, P. aeruginosa, S. aureus* and *S. epidermidis* using different molecular weight fractions (<3 kDa, 3–10 kDa, >10 kDa) and culture supernatant.

### 3.3. Impact of three selected strains (241155E, 241170E and 241174E) on biofilm formation of *K. oxytoca* and *S. epidermidis* in microfluidic flow-cells

Based on the initial screen for static biofilm inhibition using the crystal violet assay (Fig. 2, Fig. 3), three strains with high inhibitory potential 241155E (*B. muris*), 241170E (*B. fragilis*) and 241174E (*B. caecimuris*)), were selected to further evaluate the biofilm inhibition in a continuous microfluidic flow cell. The microfluidic flow cell has been established for monitoring and quantifying the biofilm formation of *K. oxytoca* (Ladewig et al., 2023), and was newly established for biofilm formation of *S. epidermidis* in this project (see Materials and Methods). To evaluate the respective inhibitory impact, biofilm formation was conducted using medium continuously supplemented with cell free cell extracts generated from the selected active strains. Confocal laser scanning microscopy (CLSM) was used to analyze the biofilm formed, which allowed the measurement of biofilm thickness, volume and the visualization of the biofilm structure. The biofilm-preventing effect was calculated as a percentage value compared to the biofilm control using pure medium (100%). The effects on biofilm formation were tested for the following combinations: Cell extracts generated from strains 241155E or 241170E on *S. epidermidis*, and from strains 241155E or 241174E on *K. oxytoca*.

Under the optimized conditions, *K. oxytoca* formed a compact biofilm with a wave-like structure (Fig. 4A), achieving a mean thickness of 13 ± 3 µm and a volume of 117 ± 28 µm³ (Fig. 4B). Under the same conditions but in the presence of the 241155E cell extract (5 mg/ml), single cells adhered to the surface and developed into microcolonies (Fig. 4, A), which showed a reduced mean thickness to 4 ± 1 µm, and a decreased significant mean volume of 47 ± 17 µm³ (Fig. 4B). This corresponds to 27% of the thickness and 40% of the volume of the medium control. Similarly, in the presence of 241174E cell extract (5 mg/ml), the biofilm formed a mean thickness of 5 ± 1 µm and a mean volume of 44 ± 12 µm³ (Fig. 4B), representing 40% of the thickness and 37 % of the volume compared to the medium control. Only the formation of microcolonies was detected no compact structure like that observed in the medium control was obtained (Fig. 4A).

**Figure 4:**
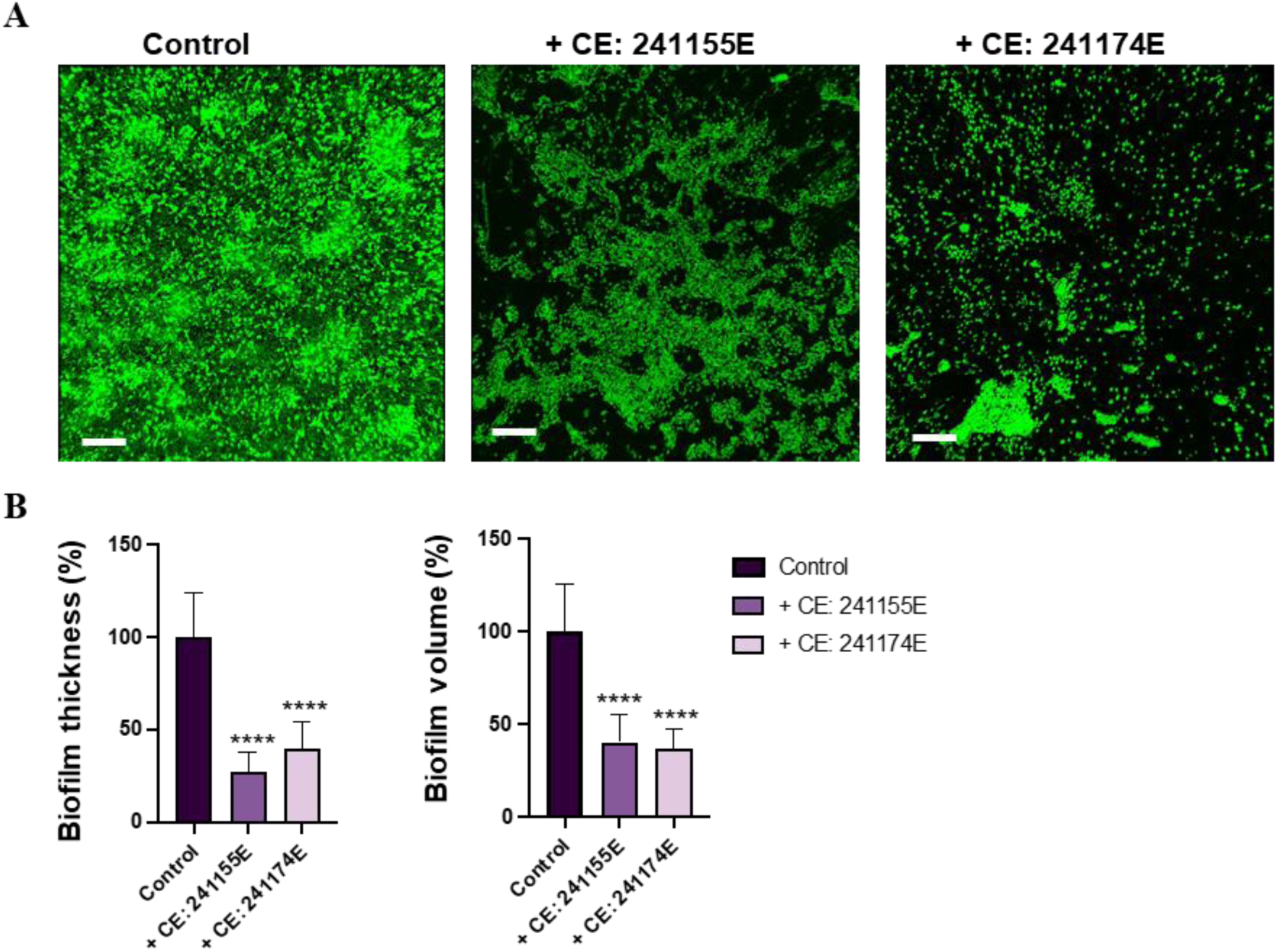
Potential of Bacteroidota isolates to inhibit biofilm formation of K. oxytoca. K. oxytoca biofilms were cultured in GC minimal media for 24 h at 30 °C, allowing bacterial cells 1 h for adhesion. The cell extracts from 241155E or 241174E were mixed with GC minimal media to generate a final concentration of 0.5%. The supplemented media was renewed every 6 h and continuously flown through the flow cell at a flow rate of 15 µL/h for 24 h at 30 °C. (A) SYTO9 was used for biofilm staining and visualized using confocal laser scanning microscopy. Representative CLSM images with scale bars representing 40 µm. (B) Analysis was performed using Zen Black (version 14.0.22.201) software and Imaris (version 9.9.0). The biofilm-preventing effect was calculated as a percentage value compared to the biofilm control (100%). Biofilm characteristics are presented in a bar plot, representing the means of four biological replicates, each consisting of four technical replicates, along with the respective standard error of the mean. Unpaired t-test was performed with GraphPad Prism 6 software with differences *p < 0.05, **p < 0.01, ***p < 0.001, ****p < 0.001 considered significant.

The optimized biofilm formation of *S. epidermidis* was established with an initial cell concentration of 3.2 × 10^5^ cells/channel and a flow rate of 18 µL/h for 20 h at 37°C. Under those conditions, *S. epidermidis* formed a planar and compact biofilm (Fig. 5A) with a thickness of 12 ± 3 µm and a volume of 224 ± 67 µm³ (Fig. 5B). Same conditions but in the presence of the 241155E cell extract (5 mg/ml), the biofilm thickness was reduced to 5 ± 1 µm, and the volume decreased to 21 ± 10 µm³ (Fig. 5, B). This represents 37 % of the thickness and 15 % of the volume of the medium control biofilm. Similarly, when exposed to the 241170E cell extract (5 mg/ml), the biofilm thickness was reduced to 4 ± 1 µm, and the volume was lowered to 34 ± 21 µm³ (Fig. 5B). These values account for 41% of the thickness and 9% of the volume compared to the medium control. In the presence of 241155E and 241170E, the surface was coated by attached single cells forming microcolonies (Fig. 5A).

**Figure 5:**
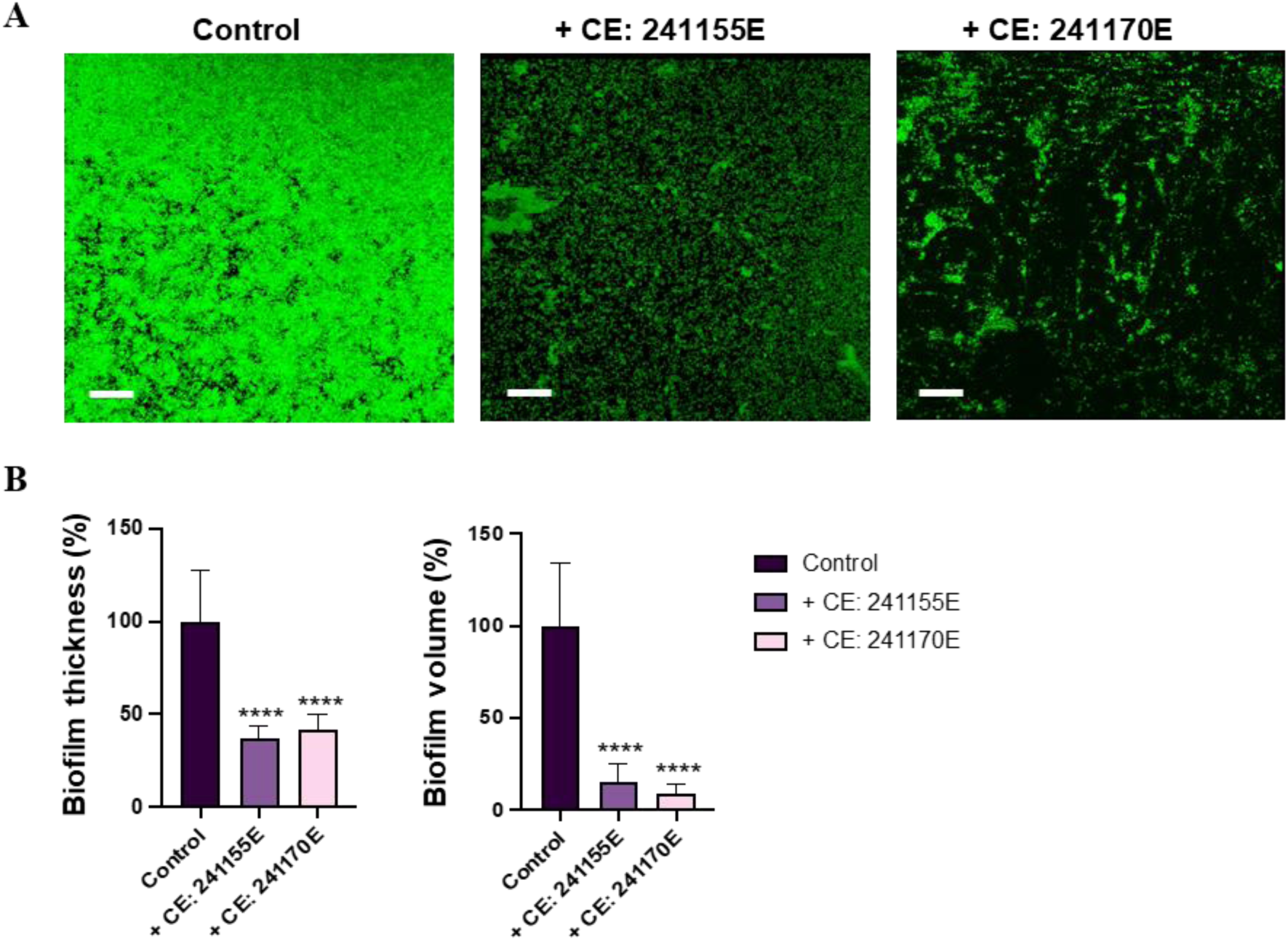
Effective inhibition of *S. epidermidis* biofilm formation by 241155E and 241170E. *S. epidermidis* biofilms were generated in Caso broth media for 20 h at 37 °C, with bacterial cells allowed 1 h for adhesion. Cell extracts from 241155E or 241170E were then mixed with Caso broth to attain a final concentration of 0.5%. This mixture was renewed every 6 h and continuously flown through the flow cell at a rate of 18 µL/h for 20 h at 37 °C. (**A**) Biofilm staining was performed using SYTO9 and visualized using confocal laser scanning microscopy, yielding representative CLSM images with scale bars set at 40 µm. (**B**) Biofilm analysis was conducted using Zen Black (version 14.0.22.201) software and Imaris (version 9.9.0). The biofilm-preventing effect was calculated as a percentage value compared to the biofilm control (100%). The biofilm characteristics were presented in a bar plot, indicating the means of four biological replicates, each comprising four technical replicates, along with their respective standard errors of the mean. Statistical analysis was carried out using an unpaired t-test with GraphPad Prism 6 software, where differences with *p < 0.05, **p < 0.01, ***p < 0.001, ****p < 0.001 were deemed significant.

Consequently, this demonstrated that the cell extracts 241155E and 241174E significantly inhibit biofilm formation of *K. oxytoca*, and the extracts 241155E and 241170E similarly inhibit biofilm formation of *S. epidermidis*. These findings are consistent with the initial characterization using the crystal violet assay and suggest that these *Bacteroides* taxa have strong potential for developing strategies to control biofilm formation and mitigate the risks associated with biofilm formation of pathogens.

### 3.4. Quorum quenching by Bacteroidota strains

The quorum quenching activity was tested using *E. coli*-based reporter strains (Weiland-Bräuer et al., 2015) in the presence of specific signaling molecules. Among 42 tested Bacteroidota strains, five showed a significant AHL-quenching activity and 13 interfered with AI-2. The AHL-quenching activity was exclusively detected in the cell extract fraction, whereas AI-2 activity was mostly found in the cell extract but also in three cases in the supernatant (Table 2). Notably, strain 241155E, 241159E, 241174E and 241191E demonstrated a quenching activity for both AHL and AI-2. Among the strains with QQ activity against AI-2, five (241174E, 241160E, 241158E, 241155E and 241166E) inhibited *K. oxytoca* biofilm formation by ≥ 20%. However, some isolates, such as 241175E, showed QQ activity against AI-2 but no inhibition of *K. oxytoca* biofilm formation in the crystal violet assay (Fig. 2), suggesting that low concentrations of QQ inhibitors may not sufficiently block QS signals. Similarly, isolates showing QQ activity against AHL did not inhibit biofilm formation in *P. aeruginosa*. This might be due to the fact that *P. aeruginosa* has three quorum sensing systems, some of which are not affected by QQ activity.

**Table 2:**
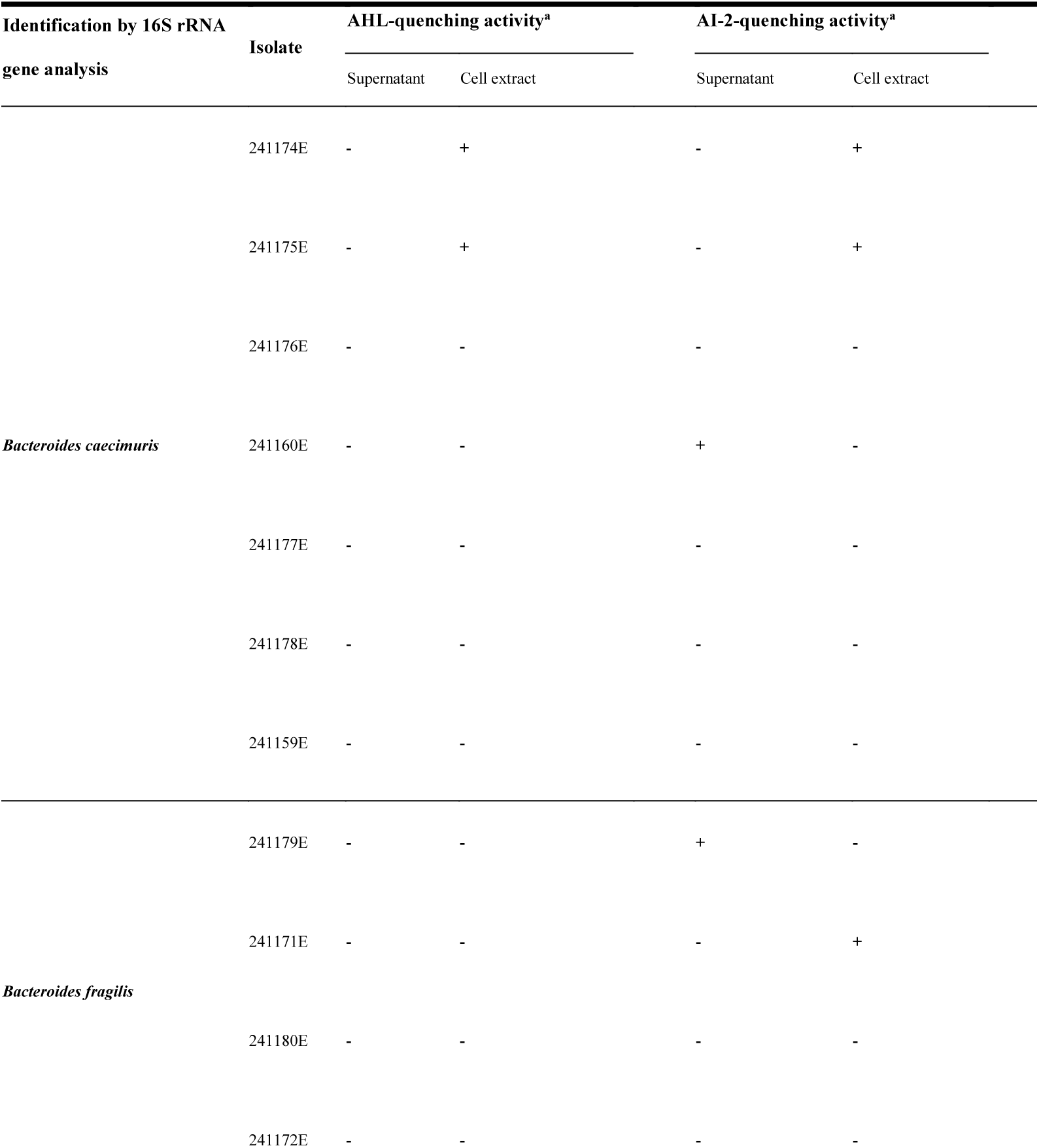
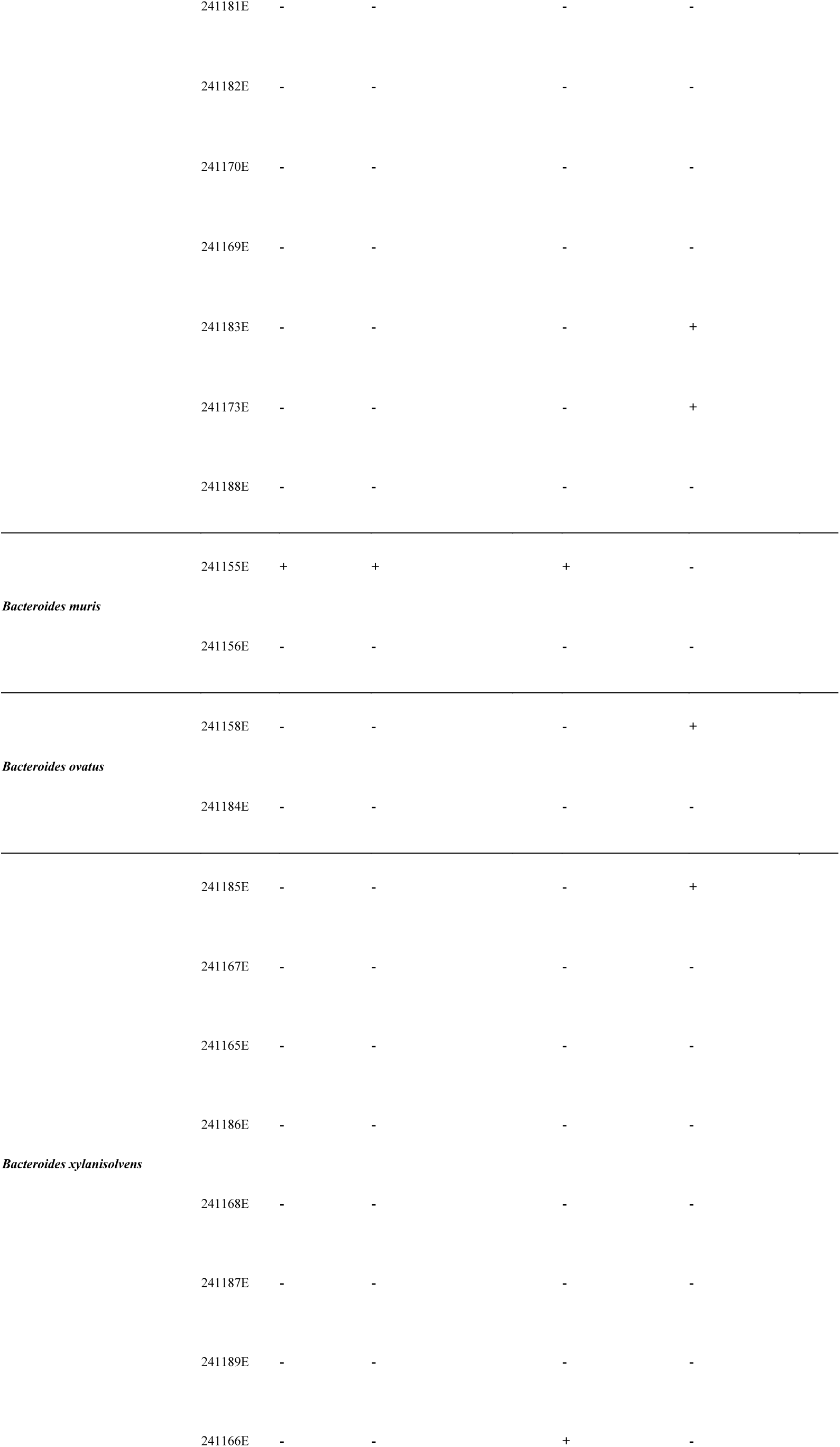
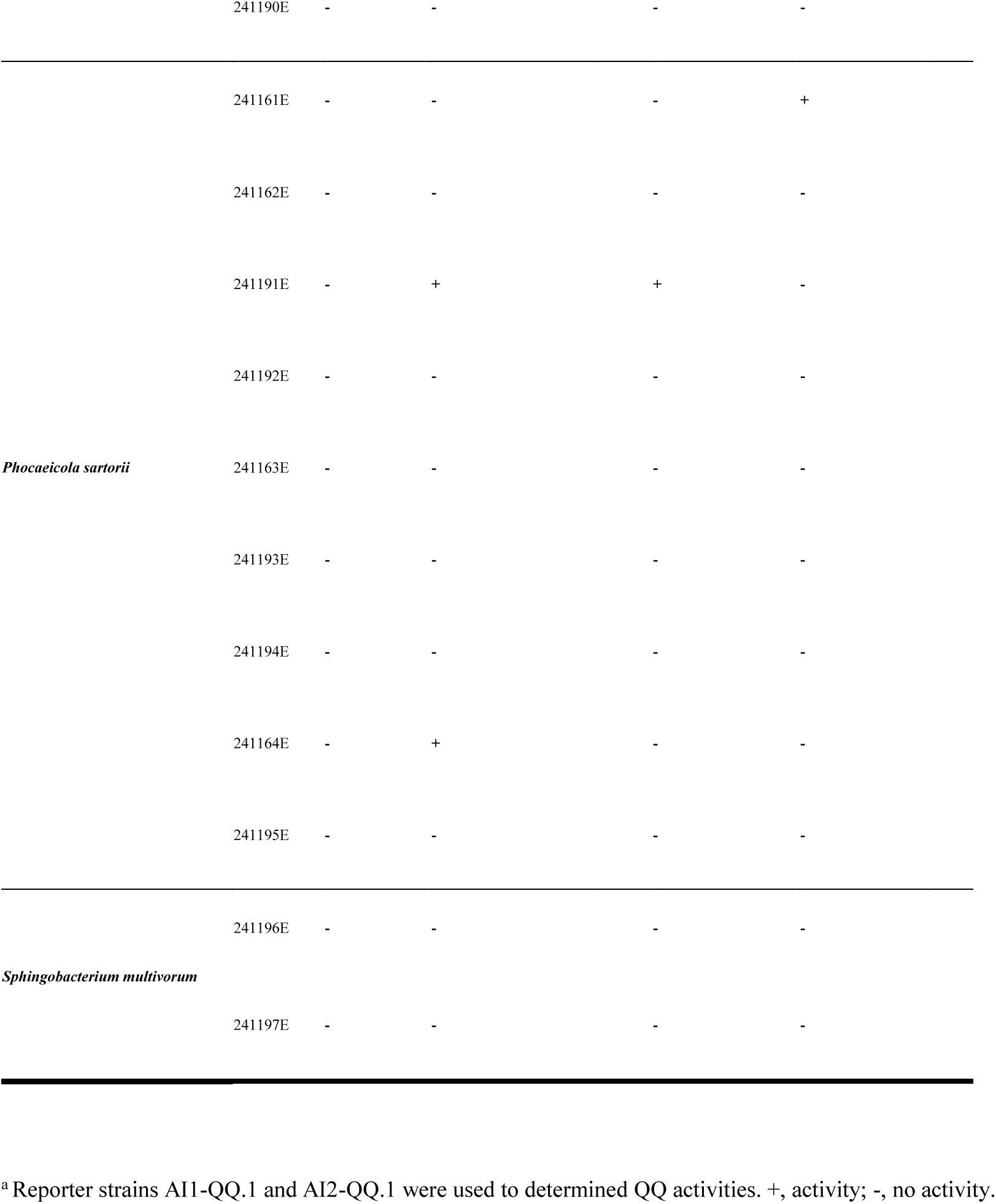
Screening for quorum-quenching-activity of the isolated Bacteroidota strains.

### 3.5. Bacteroidota strains show bile salt hydrolysis activity

Next, we tested the strains for bile salt hydrolysis activity (BSH, see Materials and Methods). BSH enzymes cleave the bond between bile acids and conjugated amino acids (glycine or taurine), producing free bile acids. This affects their antimicrobial activity, solubility, intestinal reabsorption, and gut microbiome composition. The test-medium used contained bile acids, which are deconjugated in the presence of bile salt hydrolase (Fig. 6, A). The hydrolysis of bile acids appears as a precipitation zone around the punched hole where the culture was inoculated. The diameter of all precipitation zones was measured (Fig. 6, B). Nearly all strains exhibited BSH activity, underscoring the prevalence of this enzymatic capability within the studied samples (Fig. 6, C). Only the three strains 241155E, 241158E and 241196E displayed no detectable BSH activity. The widespread BSH activity among our isolates, with only three exceptions, suggests their potential impact in bile acid metabolism within the gut microbiome.

**Figure 6:**
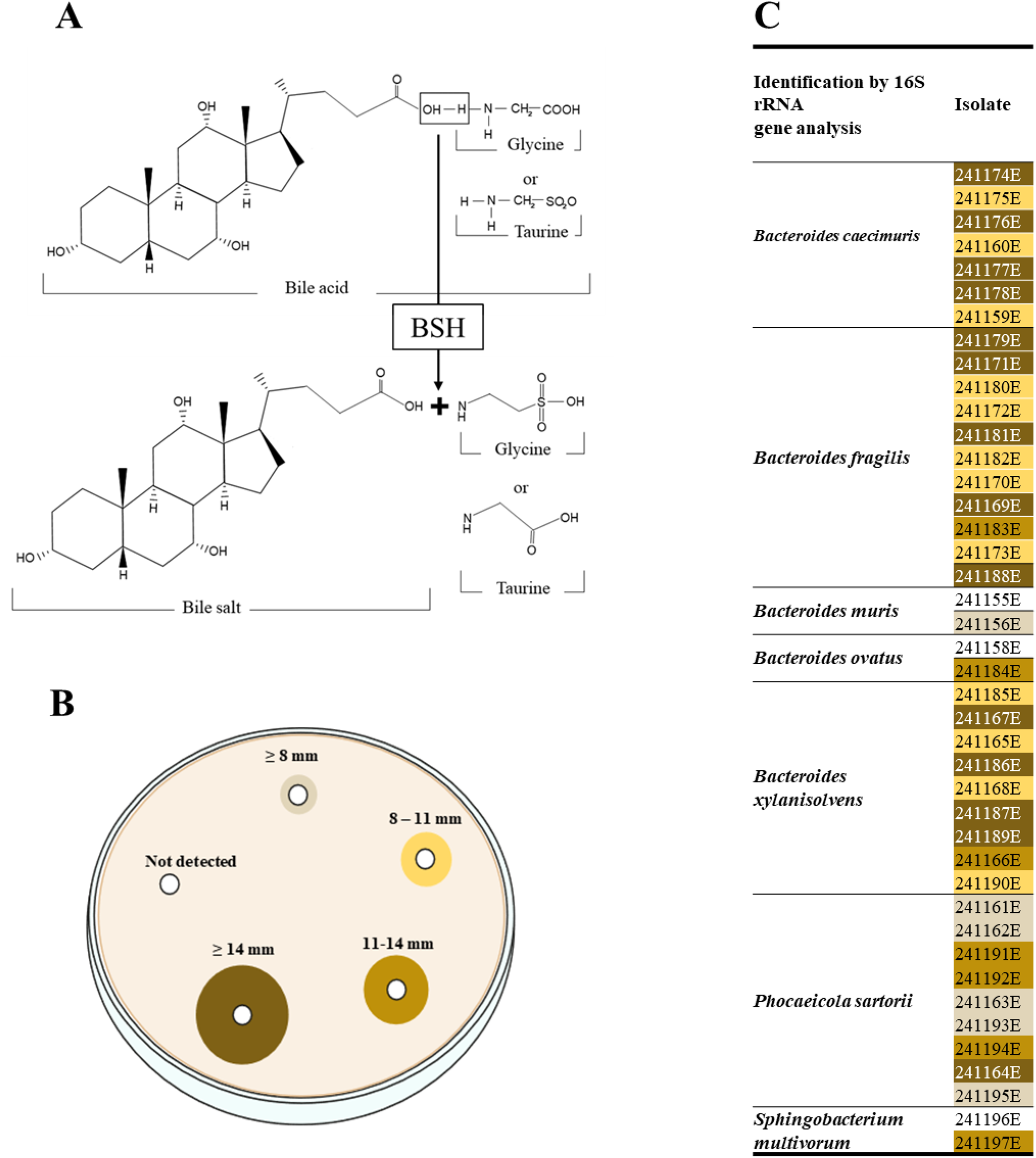
BSH activity of the isolated Bacteroidota strains. (**A**) Conjugation of bile acids with glycine or taurine results in the formation of bile salts. These bile salts are hydrolyzed by the microbial enzyme BSH, a process known as deconjugation. (**B**) BSH activity across Bacteroidota isolates is categorized by the diameter of the precipitation zone in millimeters. (**C**) Illustration of BSH activity levels, with darker shades representing higher activity. The absence of shades indicates that no BSH activity was detected for those specific isolates.

## 4. Discussion

The systematic investigation of newly isolated Bacteroidota strains uncovered a diverse repertoire of probiotic properties, including varying capacities for biofilm inhibition, QQ activity, and BSH activity, functional characteristics that strengthen their candidacy as NGPs (Fig. 7). To mention a few, strains like *B. muris (*241155E) and *B. ceacimuris* (241174E) show strong biofilm inhibition, particularly against *K. oxytoca* and *S. epidermidis* and are promising candidates for biofilm-related interventions. Strains such as *B. fragilis* (241170E), and *B. ceacimuris* strains (241174E and 241175E) offer a combination of significant biofilm inhibition and BSH activity, which could be an advantage for gut microbiota modulation and metabolic health. Moreover, the presence of QQ activity of *P. sartorii* strains (241155E, 241174E, 241175E, and 241192E) highlights their potential to disrupt pathogenic communication systems, further enhancing their probiotic efficacy.

**Figure 7:**
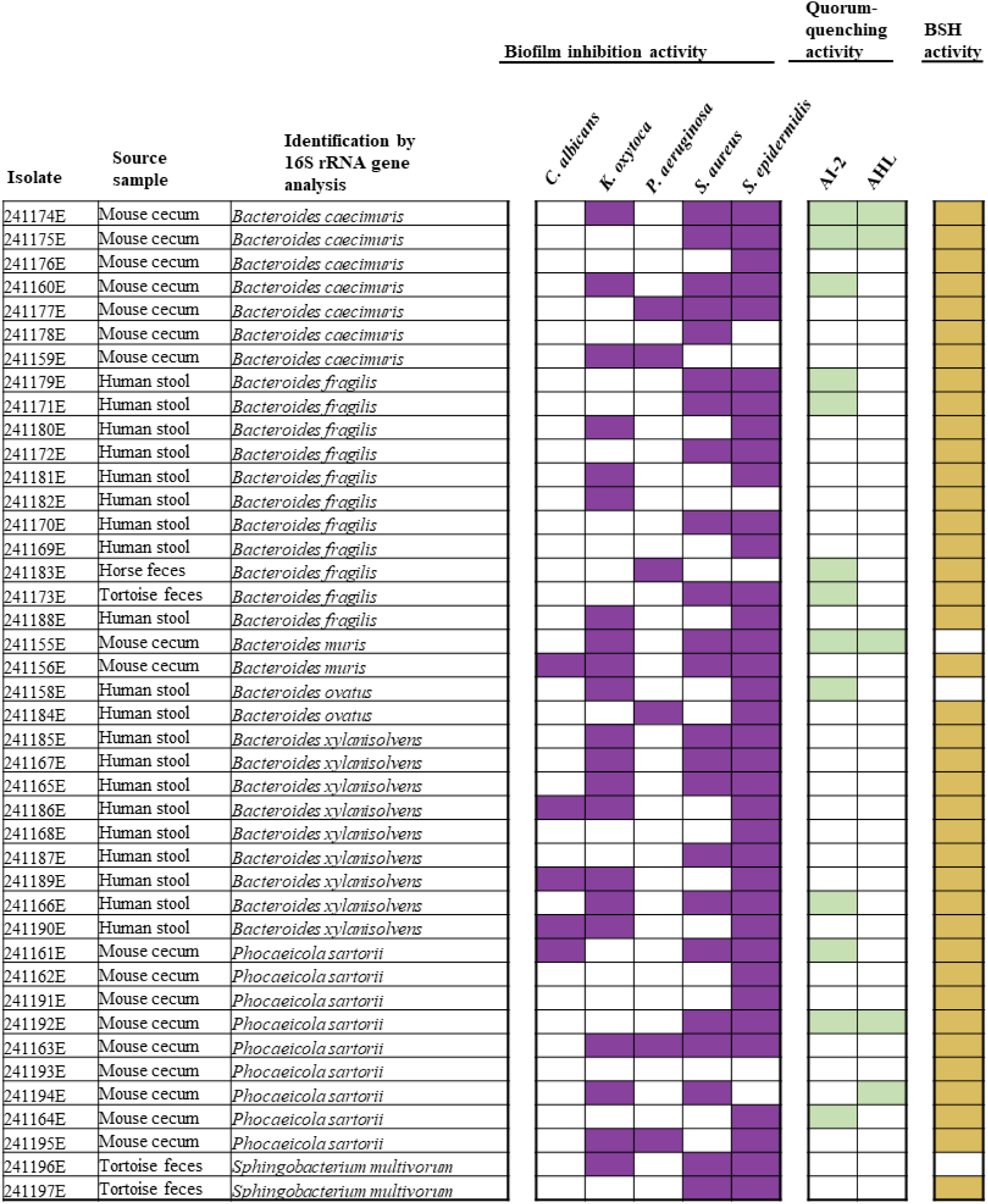
Overview of the characteristics of the isolated Bacteriodota strains. The probiotic and antimicrobial properties of the Bacteroidota isolates were tested. For the determination of biofilm-inhibiting activity, biomass was assessed using the crystal violet assay. White squares indicate < 20% inhibition, while purple squares indicate ≥ 20% inhibition. To determine the QQ activities, reporter strains AI1-QQ.1 and AI2-QQ.1 were used (Weiland-Bräuer et al. 2015). White squares indicate no QQ activity, and green squares indicate QQ activity. The BSH activities were determined by measuring the precipitation zone diameter. Ocher squares show the presence of BSH activity, and white squares indicate its absence.

Overall, among all the tested strains, *B. muris* strain 241155E stands out as the most promising candidate due to its strong biofilm inhibition, particularly against *K. oxytoca* and *S. epidermidis*, along with its significant QQ activity. This combination makes 241155E especially well-suited for disrupting pathogenic biofilms and enhancing probiotic efficacy, addressing the critical need for novel antimicrobial interventions. *B. muris* is a newly identified species within the genus *Bacteroides*, first isolated from the mouse cecum by Fokt et al. (2022). Fokt and colleagues further discovered that the two isolated strains of *B. muris* may play a part in carbohydrate metabolism and likely utilize antimicrobial peptides to affect other members of the mouse gut microbiota, pointing to potential involvement in shaping microbial interactions within the gut ecosystem, though their specific functions remain largely unexplored.

Similarly, *P. sartorii* and *B. caecimuris* are relatively newly identified species (Clavel et al., 2010; Lagkouvardos et al., 2016). Consequently, there is currently only limited information available on *P. sartorii* and *B. caecimuris* specific role(s) within the gut microbiota or its potential as a probiotic. Recent studies have mainly focused on the genetic and phenotypic characterization of *P. sartorii* and *B. caecimuris*, but their impact on gut health and interactions with other microbial species have yet to be thoroughly explored.

*B. fragilis* is a key member of the human gut microbiota, constituting about 1-2% of cultured fecal bacteria (Lee et al., 2024; Sears et al., 2014). The strain *B. fragilis* ZY-312, isolated from human feces of a healthy person, has already been identified as a potential candidate for NGP (Wang et al., 2017). Notably, this strain has demonstrated the ability to enhance macrophage phagocytic activity, promoting their polarization toward the M1 phenotype. This finding highlights its potential role in modulating immune responses and combating infections (Deng et al., 2016; Sun et al., 2019). The immunomodulatory effects exhibited by this promising strain merit further investigation to elucidate its potential applications in the treatment of diverse inflammatory and infectious diseases.

In general, understanding these microbial interactions within the gut microbiome is essential, as the gut microbiota plays a crucial role in digestion, immune system support, nutrient absorption, protection against harmful microorganisms, and stabilization of the intestinal barrier (McCallum & Tropini, 2024). Nonetheless, within this complex community, biofilms create a protective environment for microorganisms in the gastrointestinal tract, allowing them to evade the immune system and resist antibiotic treatments (McCallum & Tropini, 2024; Motta et al., 2021). Consequently, biofilms of pathogenic microorganisms are responsible for nearly 60% of chronic and recurrent infections, posing a significant challenge in clinical management (Jandl et al., 2024). This protective biofilm environment can promote persistent infections, particularly in the context of inflammatory bowel diseases (IBD). In IBD, including Crohńs disease and ulcerative colitis, biofilm compromise the intestinal barrier, increasing gut permeability and triggering chronic inflammation.

The World Health Organization (WHO) has highlighted the urgent need for new antimicrobial agents to address critically important pathogens like *Klebsiella, Staphylococcus* and *Pseudomonas* species, which are developing resistance to last-resort antibiotics (Garvey, 2023; Mancuso et al., 2021; WHO, 2024). Pathogenic biofilms in the gut, such as those formed by *P. aeruginosa*, *S. aureus* and *C. albican*s, can interact with the mucosal immune system, exacerbating immune responses and promoting a circle of inflammation and microbial persistence (Srivastava et al., 2017). Consequently, targeting biofilm formation could reduce infection severity and inflammation, making it a promising focus on IBD treatment research (Srivastava et al., 2017). In this study, five Bacteroidota isolates were identified that inhibit *C. albicans* biofilm formation, six isolates inhibit *P. aeruginosa* biofilm formation, and 25 biofilm formation of *S. aureus*.

Quorum sensing (QS) interference has emerged as a promising therapeutic strategy against bacterial infections, particularly through the use of quorum quenching (QQ) molecules (Vashistha et al., 2023). While QQ molecules are found across both prokaryotes and eukaryotes, as reviewed in Fetzner (2015), the QQ capabilities of Bacteroidota remain largely unexplored, highlighting the importance of our finding that some Bacteroidota show QQ activity (Table 2). Our study investigated QQ activity in Bacteroidota across multiple host sources. The inclusion of non-human hosts in our study underscores the importance of exploring diverse microbiomes for discovering novel antimicrobial properties that may not be present in human-derived strains alone. Notably, we identified 14 isolates with QQ activity, with only 4 originating from human stool samples. This finding further highlights a significant knowledge gap in understanding QQ mechanisms within Bacteroidota. The limited knowledge and research on Bacteroidota’s QQ potential may be attributed to several factors. Only a few studies have systematically explored non-human hosts, some strains were only recently discovered, and many remain inadequately characterized. Nevertheless, our findings indicate significant potential for future investigations into the quorum quenching capabilities of Bacteriodota, particularly in developing targeted anti-biofilm approaches that could offer advantages over conventional antibiotics by preserving beneficial gut microbiota. The development of such selective therapies could potentially revolutionize the treatment landscape for gastrointestinal disorders by specifically targeting pathogenic biofilms whilst maintaining microbial homeostasis.

These findings align with research showing that examining microbiota from varied hosts can uncover unique microbial properties with potential valuable applications. For instance, in a recent study by Kim et al. (2018), gut microbial diversity was examined in weaning swine fed probiotic bacteria that inhibit quorum sensing molecules. *Lactobacillus acidophilus* strain 30SC, isolated from pig intestine, effectively inhibited autoinducer-2 (AI-2) activity in entero-hemorrhagic *E. coli* O157:H7, without compromising bacterial viability. When administered to weaning swine, this probiotic intervention resulted in a selective modulation of gut microbiota, characterized by a reduction in coliform populations and an increase in beneficial lactobacilli, while maintaining stable levels of total aerobes and yeasts. These findings suggest that quorum-quenching probiotics can beneficially alter the gut microbiota in swine, enhancing their gastrointestinal health and performance (Kim et al., 2018). Similarly, recent research involving two AHL-degrading bacterial enrichment cultures derived from shrimp gut microbiota demonstrated their potential in improving gut health in marine fish larvae. Two enrichment cultures, EC3 and EC5, successfully colonized the gut of marine fish larvae, with EC5 demonstrating particular efficacy in enhancing larval survival when challenged with AHL molecules. EC5’s ability to neutralize AHL-mediated bacterial virulence highlights the potential of quorum sensing interference as an alternative to conventional antibiotics in marine aquaculture (Tinh et al., 2008). Thus, the systematic exploration of microbiota across diverse host environments emerges as a promising strategy for discovering novel therapeutic interventions.

### Implications and Future Directions

The functional diversity observed in our Bacteroidota isolates, particularly their quorum quenching and biofilm inhibition capabilities, positions them as promising candidates for probiotic applications. The BSH activity further supports their viability as probiotics, indicating their ability to survive in the gut environment. Furthermore, the mouse cecum is apparently a promising environment to detect QQ activity of Bacteroidota, since the majority of the effective strains originate from those samples (Fig. 7). Its high microbial competition and complex ecosystem might foster strains with strong biofilm inhibition, QQ activity, and BSH function. Similar complex ecosystems may also yield promising probiotic candidates for future development of therapeutic strains.

Despite their potential benefits, the application of Bacteroidota as probiotics presents several significant challenges, primarily due to regulatory, safety, and biological complexities. In addition, their complex patterns of evolution create hurdles in their characterization and adequate application in industries and dairy products (Ranjan et al., 2022). These challenges are compounded by the unique characteristics of Bacteroidota, a phylum known for its variability in pro-inflammatory surface antigens and its role in gut commensalism and opportunism. Firstly, the global regulatory landscape for probiotics is inconsistent, which complicates the development and marketing of probiotic products, including Bacteroidota. This inconsistency affects consumers, healthcare professionals, and industry stakeholders, as there is a lack of uniform guidelines and labeling standards that ensure safety and efficacy (Szajewska & Vinderola, 2024). The need for harmonized regulations is critical to facilitate the safe and effective use of probiotics, including those derived from Bacteroidota. Safety concerns are particularly pronounced with Bacteroidota due to their dual role as commensal and opportunistic bacteria. The safety of probiotics is a paramount concern, as highlighted in the context of gastrointestinal diseases, where stress resistance, post colonization quantification, and evaluation models are critical for assessing probiotic efficacy and safety (Wolfe et al., 2023).

Hudson and Egan (2022) have highlighted the need for a comprehensive understanding of virulence factors of Bacteriodota, particularly in marine environments where these bacteria could pose a threat to marine eukaryotes. Additionally, the integration into food production systems requires thorough investigation to ensure their safety and efficacy (Awad et al., 2022; Venugopalan et al., 2010). Future research should focus on key areas to expand our understanding of the probiotic potential of these isolates. In this respect i*n vivo* studies are essential to confirm their safety and efficacy as next-generation probiotics. Investigation of the molecular mechanisms is crucial for advancing microbial control strategies like prevention or inhibition. Additionally, investigating how Bacteroidota isolates influence gene expression related to biofilm formation could uncover new biofilm control targets. Genomic analysis, including whole-genome sequencing and pangenomic analysis can further identify genetic factors contributing to the isolates’ functional diversity, supporting more targeted probiotic applications (Apostolakos et al., 2022).

## 5. Conclusion

This study highlights the rich functional diversity and probiotic potential of Bacteroidota isolates from human and mouse samples. The observed quorum quenching activity, biofilm inhibition capabilities, and bile salt hydrolase activity collectively suggest that these strains may play significant roles in modulating microbial communities and promoting host health. Our findings contribute to the growing body of knowledge on non-traditional probiotic candidates and open new avenues for developing next-generation probiotics targeting specific health outcomes. These findings can further be validated by genome analysis and identifying the respective genes for probiotic properties, which will aid in developing a prediction tool based on the core genome analysis.

## Acknowledgments

This research conducted with the financial support funded by Bundesministerium für Bildung und Forschung (BMBF) (031B0846D to R. Schmitz-Streit, within the BaPro consortium) and the German Science foundation (DFG) as part of the CRC1182 “Origin and function of metaorganisms” (Z2 project, Schmitz-Streit). We particularly thank Nicole Pinnow for technical support and Dr. Corinna Bang (IKMB, Kiel) for providing the human feces samples.

The authors declare no conflict of interest.

## Supplementary Materials

**Figure S1:** Design and 3D-printed wash assist device. (A) The schematic design, created using Autodesk Inventor (Munich, Germany) and Prusa Slicer (Prague, Czech Republic), features engineered pins and empty channels for optimal liquid flow and minimal biofilm disturbance during wash steps. (B) Actual 3D-printed device made from polyethylene terephthalate glycol on a Prusa MK3S printer.

