## Supplement Figure 1 for "Exploring new Bacteroidota strains: Functional Diversity and Probiotic Characteristics"

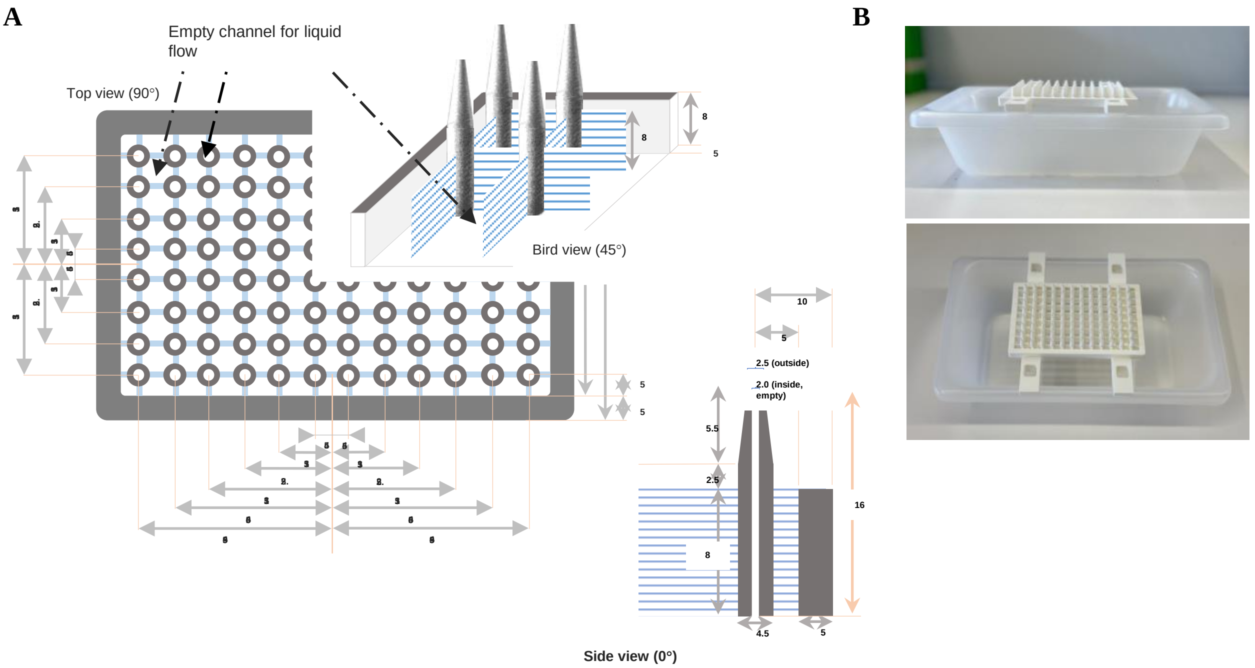


**Figure S1: Design and 3D-printed wash assist device. (A)** The schematic design, created using Autodesk Inventor (Munich, Germany) and Prusa Slicer (Prague, Czech Republic), features engineered pins and empty channels for optimal liquid flow and minimal biofilm disturbance during wash steps. (**B)** Actual 3D-printed device made from polyethylene terephthalate glycol on a Prusa MK3S printer.
